# Expression of the endocannabinoid system in the human airway epithelial cells – Impact of sex and chronic respiratory disease status

**DOI:** 10.1101/2020.03.01.971960

**Authors:** Matthew F. Fantauzzi, Jennifer A. Aguiar, Benjamin J-M. Tremblay, Toyoshi Yanagihara, Abiram Chandiramohan, Spencer Revill, Min Hyung Ryu, Chris Carlsten, Kjetil Ask, Martin Stämpfli, Andrew C. Doxey, Jeremy A. Hirota

**Affiliations:** Firestone Institute for Respiratory Health – Division of Respirology, Department of Medicine, McMaster University, Hamilton, ON, L8N 4A6; McMaster Immunology Research Centre, McMaster University, Hamilton, ON, Canada, L8S 4K1; Department of Biology, University of Waterloo, Waterloo, ON, Canada N2L 3G1; Division of Respiratory Medicine, Department of Medicine, University of British Columbia, Vancouver, BC, Canada, V6H 3Z6

## Abstract

Recreational and medicinal cannabis consumption in the past 12 months has been reported in 1/5^th^ of Canadians, with greater use in males relative to females. Cannabis smoking is the dominant route of delivery in consumers, with the airway epithelium functioning as the site of first contact for inhaled phytocannabinoids. The endocannabinoid system is responsible for mediating the physiological effects of inhaled phytocannabinoids. Acute cannabis smoke inhalation can result in bronchodilation, which may have applications in chronic respiratory disease management. In contrast, chronic cannabis smoke inhalation is associated with reduced lung function and bronchitis, which challenges potential applications in the lung. The contribution of the endocannabinoid system in the airway epithelium to either beneficial or harmful physiological responses remains to be clearly defined in males and females and those with underlying chronic respiratory disease.

To begin to address this knowledge gap, a curated dataset of 1090 unique human bronchial brushing gene expression profiles was created from Gene Expression Omnibus deposited microarray datasets. The dataset included 616 healthy subjects, 136 subjects with asthma, and 338 subjects with COPD. A 27-gene endocannabinoid signature was analyzed across all samples with sex and disease specific-analyses performed. Immunohistochemistry and immunoblots were performed to confirm *in situ* and *in vitro* protein expression of select genes in human airway epithelial cells.

We confirm three receptors for cannabinoids, CB_1_, CB_2_, and TRPV1, are expressed at the protein level in human airway epithelial cells *in situ* and *in vitro*, justifying examining the downstream endocannabinoid pathway more extensively at the gene expression level. Sex status was associated with differential expression of 6/27 genes. In contrast, disease status was associated with differential expression of 18/27 genes in asthmatics and 22/27 genes in COPD subjects. We confirm at the protein level that TRPV1, the most differentially expressed candidate in our analyses, was up-regulated in airway epithelial cells from asthmatics relative to healthy subjects.

Our data demonstrate that endocannabinoid system is expressed in human airway epithelial cells with expression impacted by disease status and minimally by sex. The data suggest that cannabis consumers may have differential physiological responses in the respiratory mucosa, which could impact both acute and chronic effects of cannabis smoke inhalation.

## INTRODUCTION

Cannabis is the most commonly consumed illicit drug worldwide, with prevalence expected to increase due to trends in legalization and innovations in medicinal applications(1). Canada legalized cannabis in 2017 and concomitantly initiated the annual Canadian Cannabis Survey to monitor the perception and use patterns, demonstrating 20% of the general population consumes cannabis(2-4). Over 90% of cannabis consumers identified combusted smoke inhalation as a route of delivery, highlighting the lung as the dominant target for cannabis exposures. Within the cannabis consumer population, there is a skewed distribution of use patterns with males more frequently to consume in the past 12 months relative to females (26% vs 18%) with a subset (12%) of the population using cannabis for medicinal purposes. The prospective design of the Canadian Cannabis Surveys has demonstrated that cannabis use over time is widely accepted at a population level for recreational and medicinal purposes with no signal for decline in use post-legalization(2-4). Additionally, the data demonstrate a strong preference for inhalation routes of delivery and a skewing between male and female use. Therefore, a generalized understanding of how a cannabis consumer responds to smoke exposure is likely to be insufficient and should include both females and males in analyses and consider both healthy individuals and those that may have underlying medical needs.

The Canadian Cannabis Surveys report of inhalation as a dominant route of delivery confirms the importance of focusing on how the lungs respond to cannabis exposures. Early clinical exposure studies demonstrated that acute cannabis smoke exposure in healthy subjects is able to provide a sustained increase in lung function, which contrasted tobacco smoke inhalation (5, 6). In asthmatic subjects, cannabis smoking is able to produce a rapid reversal of exercise-induced bronchoconstriction and minimize bronchoconstriction induced by methacholine inhalation(7, 8). Despite these objective benefits in a controlled laboratory setting, critical consideration needs to be given to the observed negative impacts on lung function in the context of increased frequency and intensity of cannabis smoking. Contrasting acute cannabis exposure, cannabis smoking over a two-month period was associated with a decrease in airway compliance that was correlated with quantity of cannabis consumed(9). Further consolidating the negative impacts of chronic cannabis use on lung health, population level analyses reveal that greater intensity of cannabis smoking is correlated with reduced lung function and increased risk of developing chronic obstructive pulmonary disease (COPD)(10, 11). Furthermore, in multiple independent cohorts, cannabis smoking has been associated with a pro-inflammatory phenotype in the lung, associated with bronchitis and impaired immune cell function (12-16). The mechanism(s) responsible for the clinical observations resulting from cannabis inhalation, whether beneficial acute or detrimental chronic exposures, remain elusive, but are likely to be influenced by the endocannabinoid system(17-19).

The endocannabinoid system is responsible for mediating the pharmacological effects of the endogenous cannabinoids anandamide and 2-arachidonylglycerol (2-AG) and phytocannabinoids present in cannabis(20-22). The first identified cannabinoid receptors were CB_1_ and CB_2_, both G-protein coupled receptors that modulate downstream cyclic-AMP signaling by inhibiting adenylyl cyclase activity(23, 24). The mechanism of acute phytocannabinoid-induced bronchodilation is suggested to be mediated via CB_1_ receptors regulating neural control of airway tone(17, 18), which may have benefits for management of bronchoconstriction in asthma or COPD. In addition to CB_1_ and CB_2_, the transient receptor potential vanilloid-1 (TRPV1) and the de-orphanized receptor GPR55 have been identified as receptors for cannabinoids(25, 26). Although the endocannabinoid signaling pathway is present throughout multiple tissues and organ systems, the lungs are of particular interest as they are the organ system targeted by cannabis inhalation. Airway epithelial cells play a critical role in lung health by acting as the first line of defence against pathogens and inhaled insults(27-29). Airway epithelial cells carry out a number of functions such as providing a physical barrier against microbial infiltration, maintaining the inflammatory microenvironment, and releasing immune mediators to recruit leukocytes to the site of insult. It has been demonstrated that the inhalation of air pollution, tobacco smoke, and cannabis smoke can compromise airway epithelial function(30-32). Notably, recent findings show that cannabis smoke exposure can lead to impaired airway epithelium barrier integrity, attenuated antiviral capacity, and exacerbated inflammatory response to immune challenges(31, 32). However, the contribution of the endocannabinoid system to these observations has not been defined. The primary cannabinoid receptors, CB_1_ and CB_2_, have been shown to be expressed in the respiratory mucosa (33, 34) and human airway epithelial cells are responsive to THC and anandamide *in vitro* (35, 36). Additional components of the endocannabinoid system including MAPK, PI3K, and protein kinase A signaling pathways downstream of receptors and enzymes responsible for cannabinoid metabolism have not been explored in an integrated fashion in human airway epithelial cells. An examination of the entire endocannabinoid system in human airway epithelial cells is required to better understand which components are dominant and likely to be functionally relevant in response to inhaled cannabis smoke. Furthermore, it remains possible that sex and disease status impact the endocannabinoid system expression, which may have functional consequences in distinct populations of cannabis consumers.

To begin out interrogation of the endocannabinoid system in human airway epithelial cells, we first generated a 27-gene endocannabinoid signature encompassing ligand recognition, signaling, and metabolism. We set out to examine if the expression of this 27-gene endocannabinoid signature was present in human airway epithelial cells and whether this was impacted by sex or disease status. The importance on examining sex and disease status on the endocannabinoid system is due to the possibility that specific populations may experience differential effects of cannabis, either positive or negative. To complete this study, we used a bioinformatic approach to analyze gene expression in 1090 unique human subject samples of airway epithelial cells isolated via bronchial brushing that included samples from males and females and individuals with asthma or COPD. We complement our bioinformatic approach with validation and confirmation of CB_1_, CB_2_, and TRPV1 in human airway epithelial cells at the protein level *in situ* and *in vitro*. Lastly, we validate a bioinformatic observation that *TRPV1* gene expression is elevated in airway epithelial cells isolated from asthmatics by performing confirmatory immunoblot analysis on primary human airway epithelial cells. Collectively, our results demonstrate that an intact endocannabinoid system is expressed in human airway epithelial cells and that disease status impacts expression to a greater extent then sex, which may have functional consequences that lead to differential responses in distinct populations of cannabis consumers.

## METHODS

### Human ethics

All studies using primary human lung material and blood were approved by Hamilton integrated Research Ethics Board or UBC Human Research Ethics Board.

### Primary Human Airway Epithelial Cells

Primary human airway epithelial cells isolated via bronchial brushings from consented healthy or asthmatic subjects were grown in PneumaCult ExPlus (Stemcell Technologies, Vancouver Canada) under submerged monolayer culture conditions and used in between passage 1 and 4. Where relevant, asthma diagnosis was confirmed with methacholine challenge and PC_20_ analysis as per ATS guidelines.

### Human Whole Lung Tissue

Non-involved tissues from lung cancer cases were used. Lungs were homogenized using a mechanical homogenizer (Omni International, Waterbuy CT), lysed in 1× lysis buffer supplemented with complete protease inhibitors (Roche), and the supernatant was collected for immunoblots.

### Human Peripheral Blood Mononuclear Cells (PBMC)

Human PBMC were isolated from the peripheral blood of healthy volunteers using Ficoll (Sigma-Aldrich) density centrifugation in Greiner LeucoSep-tubes (Sigma) according to the manufacturer’s recommendations.

### Immunohistochemistry

Formalin fixed paraffin embedded human lung tissue from healthy subjects were used for localization of CB_1_, CB_2_, and TRPV1/VR1. Three-micron thick sections were cut and stained for CB_1_ (Abcam - Ab23703 at 1:1000), CB_2_ (Abcam – Ab3561 at 1:50), and TRPV1/VR1 (Abcam – Ab3487 at 5ug/ml). All staining was performed on a Leica Bond RX system with Leica Bond reagents and heat-induced antigen retrieval in citrate buffer at pH 6. Digital slide scanning was performed using an Olympus VS120-L100 Virtual Slide System at 40X magnification with VS-ASW-L100 V2.9 software and a VC50 colour camera.

### Immunoblots

Immunoblots confirming antibody staining and protein expression in human airway epithelial cells were performed using Biorad stain free 4-20% pre-cast gradient gels and imaged on a Biorad ChemiDoc XRS+ Imaging system. For each immunoblot, 40ug of protein was added per lane. CB_1_ (Abcam - Ab23703 at 1:500), CB_2_ (Abcam – Ab3561 at 1:100), and TRPV1 (Abcam – Ab3487 at 1:500) were diluted in 5% skim milk/Tris buffered saline with 0.1% Tween-20. Primary antibody detection was performed using an anti-rabbit-HRP conjugated secondary (Cell Signaling Technology, #7074) at 1:1000 for 50 minutes at room temperature. Visualization was performed using Clarity™ Western ECL Substrate (Bio-Rad) (CB_1_ and TRPV1) and SuperSignal™ West Femto Maximum Sensitivity Substrate (ThermoFisher) (CB_2_). Total protein loading images were collected as a confirmation of equal protein loading between sample types(37).

### Gene Expression Dataset Curation, Normalization, and Statistical Analyses

Candidate genes to be analyzed were selected based on known relevance in the endocannabinoid signaling pathway. Additional genes and proteins implicated to be involved in cannabis-associated disorders and novel cannabinoid based therapeutic approaches were included and described in annotated table format (**Table 1**)(23-26, 38-56).

Public microarray experiments using Affymetrix chips (HG-U133 Plus 2, HuEx-1.0-st-v1 and HuGene-1.0-st-v1) on airway epithelial cell samples from healthy individuals or those with asthma or COPD were selected from the NCBI Gene Expression Omnibus (GEO) database. Healthy samples were further filtered by removing former or current smokers. This resulted in a total of 1090 individual samples from 27 experiments (**Table 2**) that included samples from 616 healthy subjects, 136 subjects with asthma, and 338 subjects with COPD(57-84). Within each sample population, sex was reported for a subset of samples (Healthy: 103 females/227 males, Asthma: 34 females/28 males, COPD: 48 females/93 males).

For all dataset samples, raw intensity values and annotation data were downloaded with the R statistical language (version 3.6.1; R Core Team, 2019) using the GEOquery R package (version 2.52.0)(85). Probe definition files were downloaded from the Brainarray database (version 24)(86). To obtain processed microarray gene expression values unaffected by probe CG compositional biases, the Single Channel Array Normalization (SCAN) method was used via the SCAN.UPC R package (version 2.26.0)(87) using annotation data from the Bioconductor project (version 3.9)(88). All log_2_-transformed gene expression data were unified into a single dataset, and only genes detected in all three platforms (16543) were kept for subsequent analyses. In order to reduce differential expression bias due to sex chromosome genes, only autosomal genes (15875) were included for subsequent analysis. Correction of experiment-specific batch effects was performed using the ComBat method (89) implemented in the sva R package (version 3.32.1)(90), with disease status and sex supplied as covariates. Following batch correction, all data underwent Z-Score transformation to set the mean of all samples to zero and replace expression values with a measure of variance from the mean(91). Principal component analysis (PCA) was performed using the probabilistic PCA method in the pcaMethods R package (version 1.76.0)(92). Gene expression levels were tested for significant differences via Student’s T-test with a Benjamini-Hochberg multiple testing correction using the stats R package (version 3.6.1; R Core Team, 2019). Gene expression box plots, heat maps, and PCA plots were generated with the ggplot2 R package (version 3.2.1).

## RESULTS

### Human airway epithelial cells express CB_1_, CB_2_, and TRPV1, *in situ* and *in vitro*

A curated list of genes involved in cannabinoid signaling was generated (herein called the 27-gene endocannabinoid signature) to provide a focused overview of this pathway in human airway epithelial cells (**Figure 1 and Table 1**).

**Figure 1:**
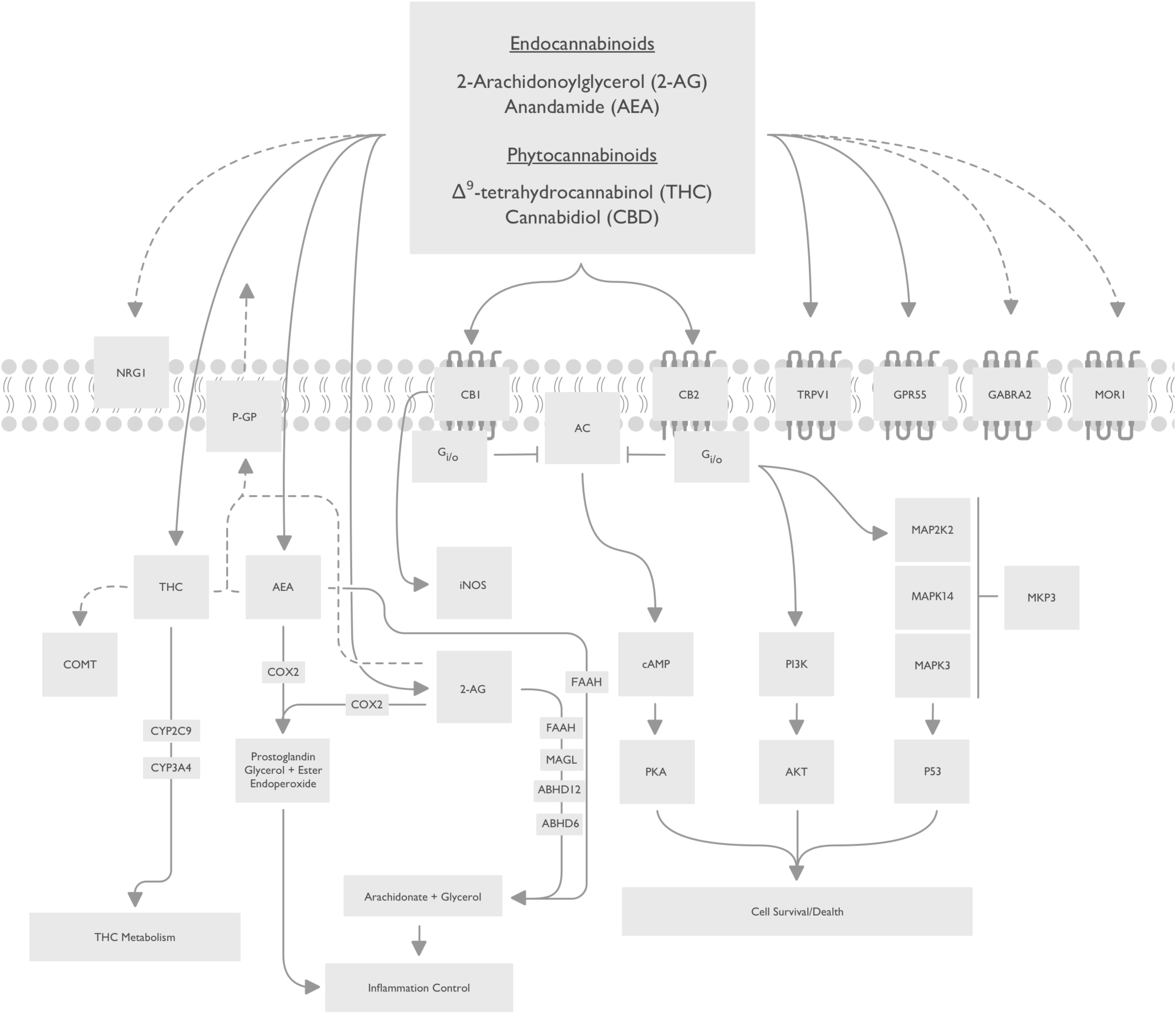
Visual representation a 27-gene endocannabinoid signature. Solid arrows indicate known relationship between candidates and ligands. Dotted arrows indicate proposed relationships. Blunted lines indicate inhibition. Candidate functions are annotated in Table 1.

To begin our characterization of the endocannabinoid system in human airway epithelial cells, we performed *in situ* localization of CB_1_, CB_2_, and TRPV1 protein in human lung tissue (**Figure 2**). We demonstrate that all three receptors are present at protein level in human airway epithelial cells *in situ* relative to negative control (**Figure 2A, C, E, G**). To validate the staining patterns observed and antibody specificity, we performed immunoblots with primary human airway epithelial cells, whole lung tissue, and PBMCs. A single band for CB_1_ was observed at approximately 45kDa in human airway epithelial cells, but not whole human lung, or PBMCs (**Figure 2B**). A dominant band for CB_2_ was observed at approximately 40kDa and accompanied by a reported 52-55kDa doublet(93) in human airway epithelial cells, with a similar pattern observed for PBMCs (**Figure 2D**). In contrast, in whole human lung the dominant band was observed at 55kDa with only a faint 40kDa band. The band patterns observed for CB_2_ are consistent with glycosylation of the N-terminus and processing of the full length peptide(93). A dominant band for TRPV1 was observed at approximately 100kDa in human airway epithelial cells and accompanied by two lower molecular weight bands at approximately 70kDa and 37kDa (**Figure 2F).** No TRPV1 bands were observed in whole human lung, while a dominant single band at 100kDa was observed in PBMCs. Total protein loading staining from a representative blot demonstrates equal loading within replicates of the same sample type and distinct protein compositions between sample types (**Figure 2H**). Collectively, our *in situ* and *in vitro* data confirm expression of CB_1_, CB_2_, and TRPV1 protein in human airway epithelial cells.

**Figure 2:**
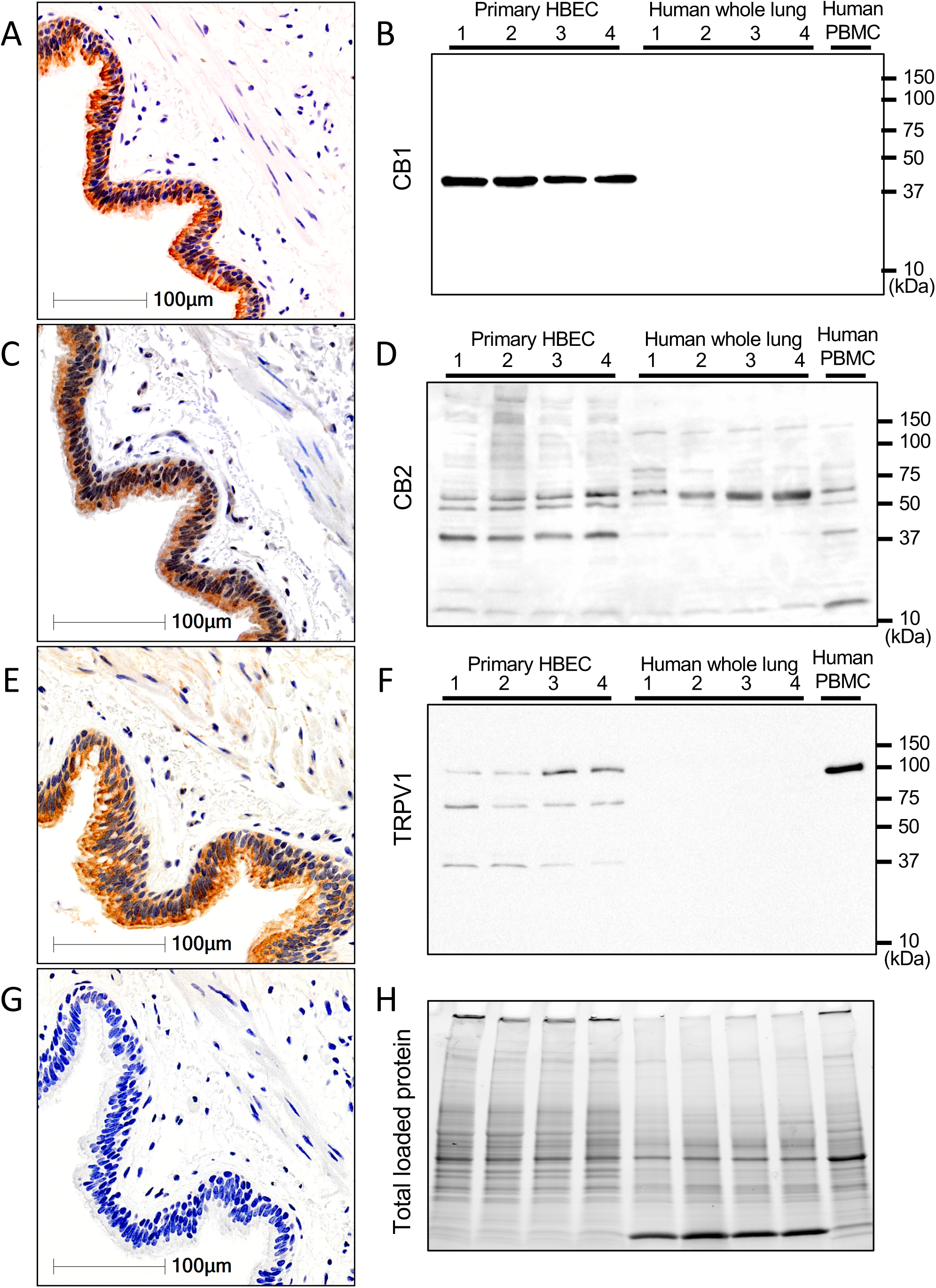
*In situ* and *in vitro* validation of CB_1_, CB_2_, and TRPV1 protein expression in human airway epithelial cells. Serial sections from a single patient donor that is representative of n=10, for immunohistochemistry of **(A)** CB_1_, **(C)** CB_2_, and **(E)** TRPV1 with **(G)** negative control. Immunoblots on primary human airway epithelial cells cultured *in vitro*, **(B)** CB_1_, **(D)** CB_2_, and **(F)** TRPV1 with **(H)** total protein loading control (n=4 airway epithelial cells, n=4 whole lung samples, n=1 PBMCs). Molecular weights (kDa) are denoted on Y-axis of immunoblots.

### Expression of endocannabinoid system genes in human airway epithelial cells from healthy male and female subjects

Sex differences in CB_1_ and CB_2_ expression levels have been reported(94), which could impact downstream responses to cannabinoid exposures. Furthermore, sex and gender differences in cannabis consumption practices have been reported(2-4). Collectively, these two factors could interact and contribute to differential responses to cannabinoids in distinct populations. To examine sex differences in the endocannabinoid system, we analyzed our 27-gene endocannabinoid signature in a curated dataset of airway epithelial cells from 616 unique healthy subjects, where the identifier of sex was available for 103 females and 227 males (**Table 2**). The 27-gene endocannabinoid signature was first analyzed in all 616 subjects to show overall trends for each gene (**Figure 3A**). Subsequently, a PCA plot was performed for all samples with sex as an identifier (**Figure 3B**). The PCA plot reveals clustering of samples from males within a larger space occupied by the female samples. Statistical analysis at the individual gene level revealed a difference in 6/27 genes(**Figure 3C**). Four genes (*ABHD6, MAPK14, NRG1*, and *PIK3CA*) were down-regulated in males relative to females, while two genes (*CYP2C9* and *GPR55*) were up-regulated in males relative to females. The gene expression patterns were overlaid on the endocannabinoid signaling pathway for qualitative visualization of global changes in the 27-gene endocannabinoid signature in males relative to females (**Figure 3D**).

**Figure 3:**
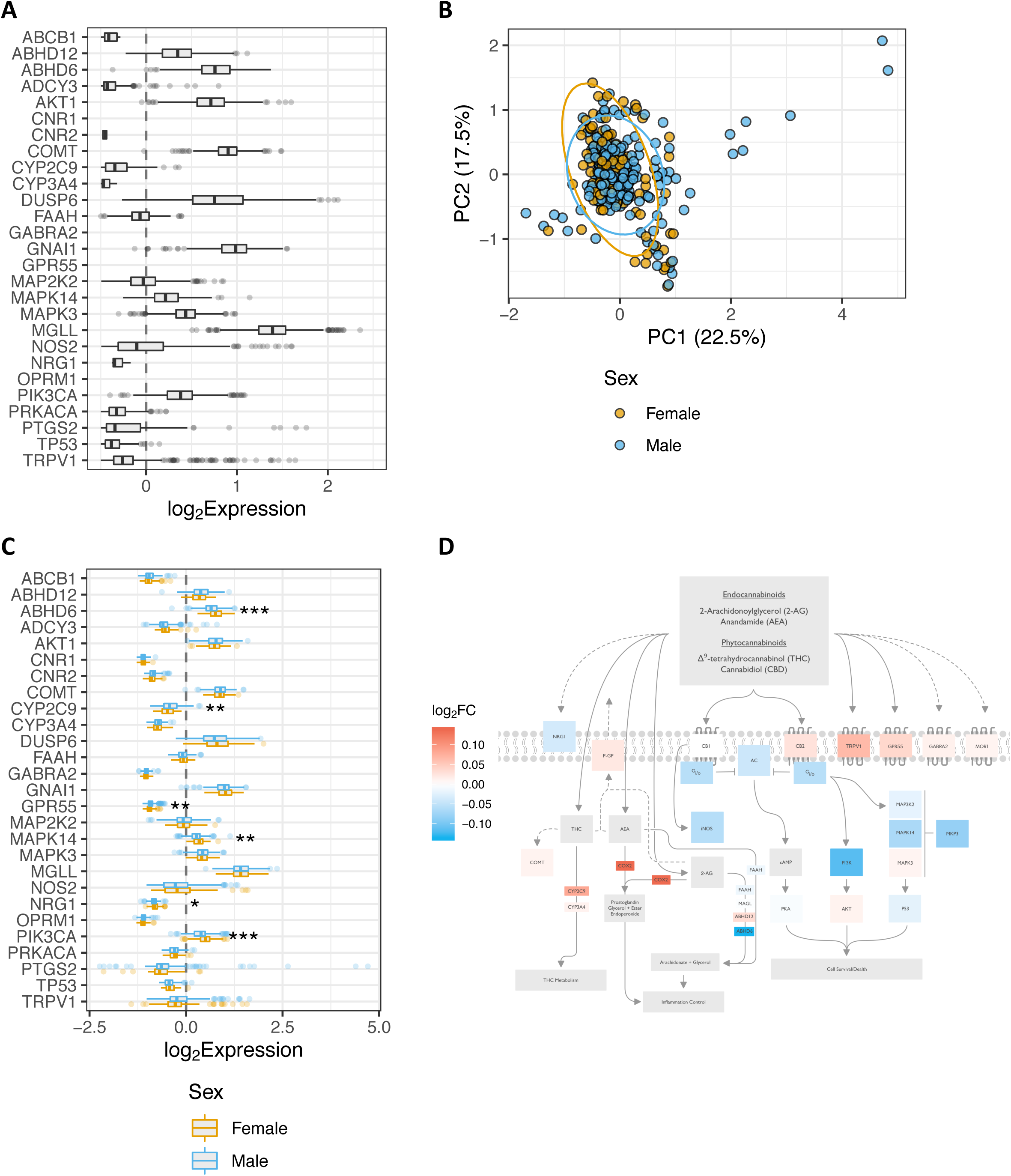
Impact of sex status on endocannabinoid system gene expression in human airway epithelial cells from healthy individuals. **(A)** Gene expression data for 616 healthy subjects with no history of smoking or chronic respiratory disease. **(B)** PCA plot of healthy females (yellow circles, n=103) and males (blue circles, n=227) generated by expression patterns of the 27-gene endocannabinoid signature. The first (22.5%) and second (17.5%) principal components were used. Ellipses were added to represent 95% confidence intervals per sex. **(C)** Healthy samples with metadata defining sex were further divided into male and female groups and plotted separately as blue and orange-outlined box plots, respectively. For both **(A)** and **(C)**, log_2_-transformed expression values were plotted as box plots. The dashed line at zero represents the global baseline of expression for the entire set of genes. **(D)** Visual representation of the differences between healthy females and males in the 27-gene endocannabinoid signature. Colour coding reflective of log_2_ fold change (log_2_FC) males relative to females. *=p<0.05; **=p<0.01; ***=p<0.001.

Collectively, our data confirm that the endocannabinoid system is expressed at gene level in human airway epithelial cells, suggesting that signaling downstream of receptors is intact, with mild sex differences observed gene endocannabinoid system gene expression.

### The endocannabinoid system is dysregulated in human airway epithelial cells from individuals with asthma and COPD

In addition to sex, disease status may also impact the expression of the endocannabinoid system in airway epithelial cells, as specific phenotypes are observed in cells isolated from asthmatics and individuals with COPD(95-97). We therefore tested the hypothesis that the 27-gene endocannabinoid signature was dysregulated in asthma and COPD.

To test this hypothesis we curated all 1090 samples that included 616 healthy subjects, 136 subjects with asthma, and 338 subjects with COPD. A PCA plot reveals clustering of samples from healthy, asthmatic, and COPD subjects, with samples from asthmatics separating from both healthy subjects and those with COPD (**Figure 4A**). Statistical analysis at the individual gene level reveals changes in 18/27 genes in asthmatics and 22/27 genes in subjects with COPD (**Figure 4B**). In asthmatics, 11/18 dysregulated genes were up-regulated (*ABCB1, ABHD6, CYP2C9, GABRA2, GNAI1, GPR55, NOS2, NRG1, PTGS2, TP53, and TRPV1*), while 7/18 were down-regulated (*ABHD12, CNR2, COMT, CYP3A4, MAP2K2, MAPK3, MGLL*). In COPD subjects, 9/22 dysregulated genes were up-regulated (*ABHD6, CNR1, CYP2C9, CYP3A4, FAAH, GABRA2, GPR55, OPMRI, TRPV1*), while 13/22 were down-regulated (*ABCB1, ABHD12, ABCY3, AKT1, DUSP6, GNAI1, MAPK14, MAPK3, MGLL, NOS2, PIK3CA, PRKACA, TP53*). The most dysregulated gene was *TRPV1*, with the largest up-regulation observed in the samples from asthmatics relative to healthy controls. The differential gene expression patterns were overlaid on the endocannabinoid signaling pathway for qualitative visualization in asthma (**Figure 4C**) and COPD (**Figure 4D**).

**Figure 4:**
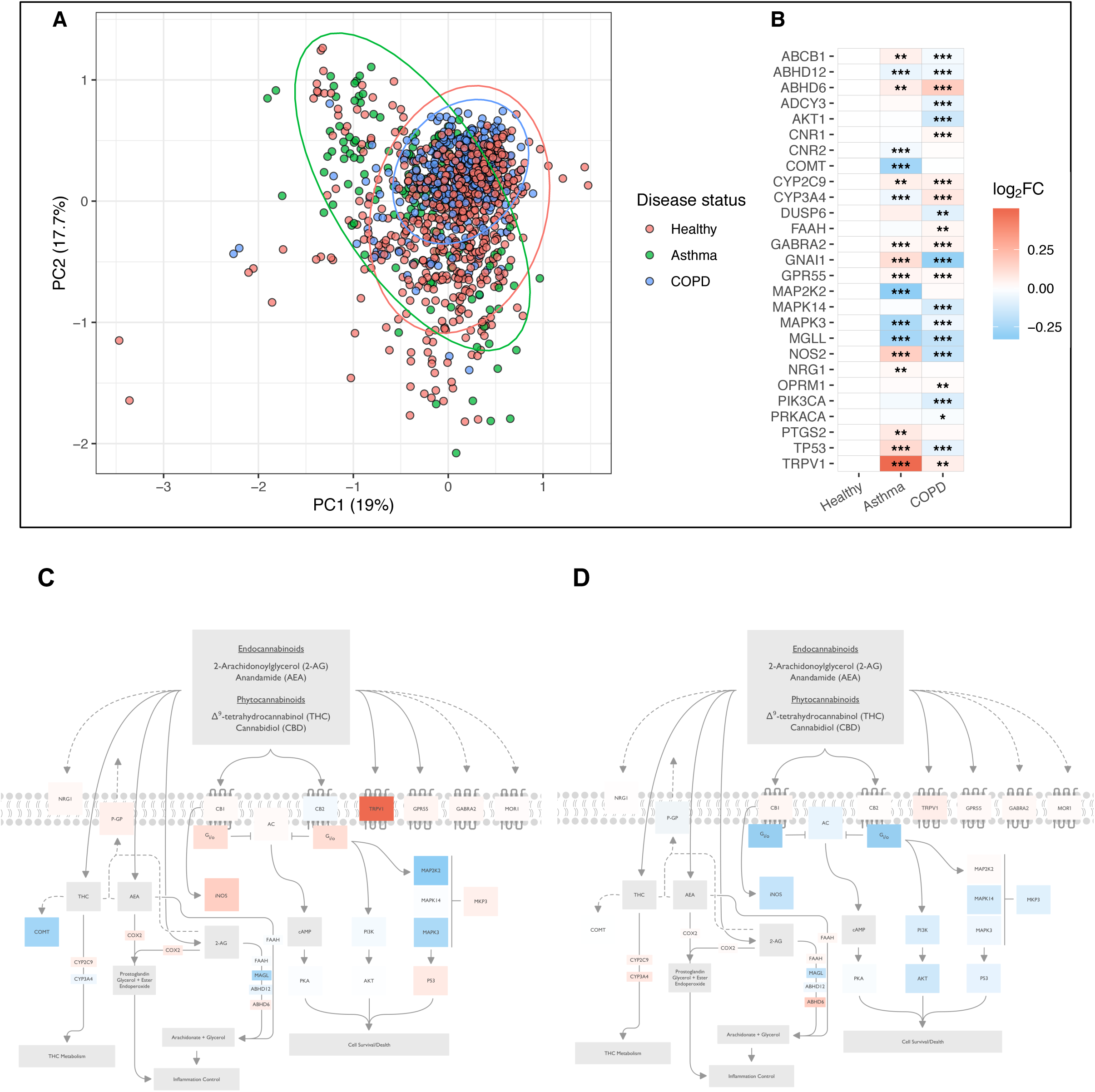
Impact of disease status on endocannabinoid system gene expression analysis in human airway epithelial cells from healthy individuals, asthmatics, and individuals with COPD. **(A)** PCA plot of healthy subjects (n=616, red circles), asthmatics (n=136, green circles) and individuals with COPD (n=338, blue circles) generated by expression patterns of the 27-gene endocannabinoid signature. The first (19%) and second (17.7%) principal components were used. Ellipses were added to represent 95% confidence intervals per sex. **(B)** Gene expression data of the 27 genes were compared between healthy, asthmatic and COPD samples. The log_2_-transformed mean expression values were compared to that of the healthy samples and shown as log_2_ fold-change (FC). Visual representation of the differences in the 27-gene endocannabinoid signature between **(C)** healthy subjects and asthmatics and **(D)** healthy subjects and individuals with COPD. Colour coding reflective of log_2_ fold chance (log_2_FC) relative to healthy subjects. *=p<0.05; **=p<0.01; ***=p<0.001.

Collectively, our data demonstrate that underlying chronic respiratory disease status is associated with a dysregulation of the endocannabinoid system at the gene level in human airway epithelial cells.

### Impact of sex on endocannabinoid system gene expression in human airway epithelial cells from asthmatics and subjects with COPD

Sex differences in incidence, age of onset, and pathology are observed in both asthma and COPD(98, 99). Cannabinoid exposures have been explored in the context of both asthma and COPD management for immunomodulatory and bronchodilation purposes (7, 8, 100). To date, the potential interaction of sex status and endocannabinoid system expression in chronic respiratory disease has not been addressed.

Taking the same approach as for healthy subjects, we analyzed a curated dataset of airway epithelial cells from 136 unique asthmatic subjects, where the identifier of sex was available for 34 females and 28 males. For COPD, we analyzed a curated dataset from 338 unique COPD subjects, where the identifier of sex was available for 48 females and 93 males (**see Table 2 for study group compositions**).

In both asthma and COPD samples, PCA plots revealed no separation between sexes with clustering of samples overlapping between disease groups (**Figure 5A, C**). At the individual gene level, no sex dependent differences were observed for any gene in the 27-gene endocannabinoid signature in either disease groups (**Figure 5B, D**).

**Figure 5:**
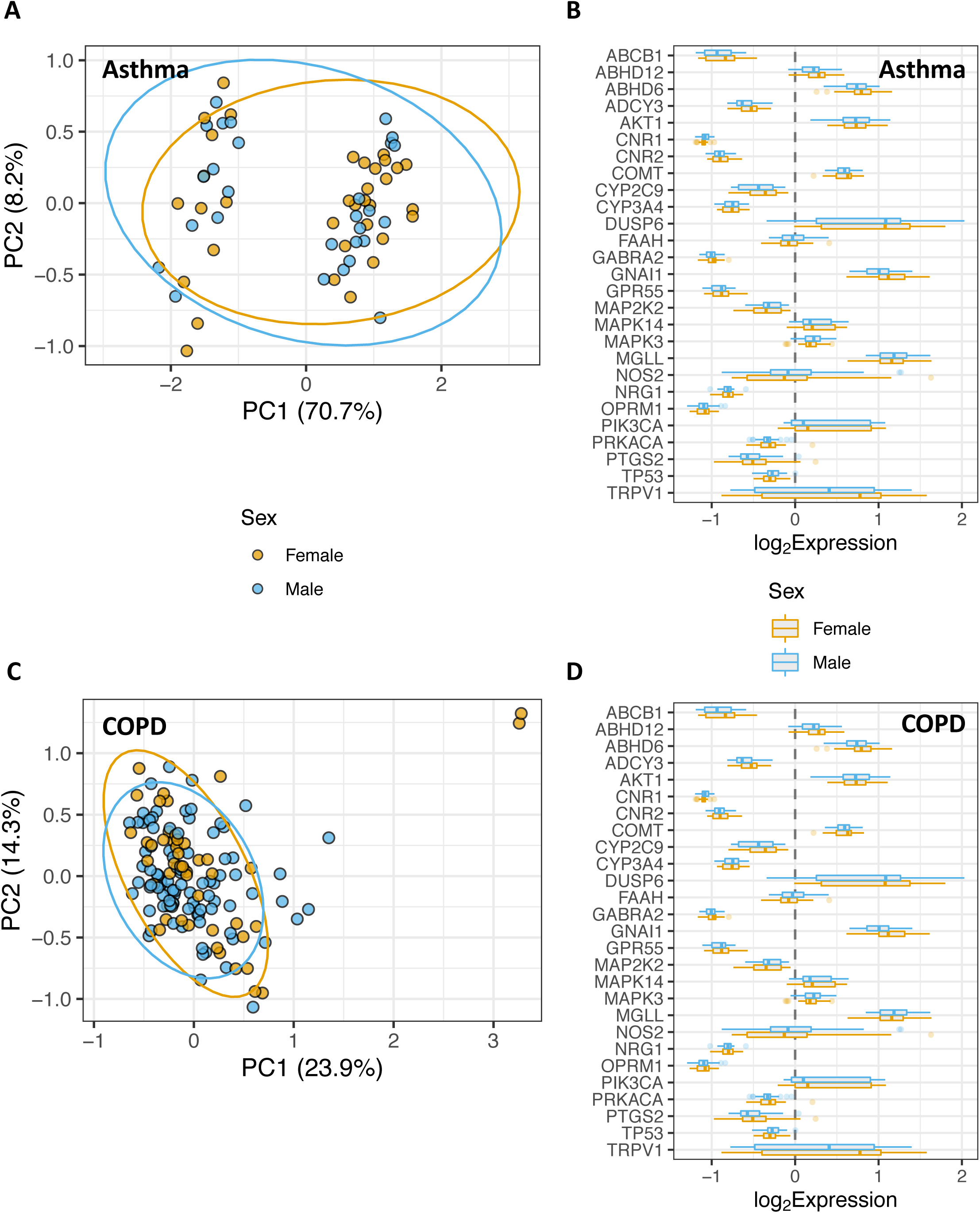
Impact of sex status on endocannabinoid system gene expression in human airway epithelial cells from individuals with chronic respiratory disease. **(A)** PCA plot of asthmatic females (yellow circles, n=34) and males (blue circles - n=28) generated by expression patterns of the 27-gene endocannabinoid signature. The first (70.7%) and second (8.2%) principal components were used. **(B)** Asthmatic samples divided into female and male, and plotted separately as blue and orange-outlined box plots, respectively. **(C)** PCA plot of females (yellow circles, n=48) and males (blue circles - n=93) with COPD generated by expression patterns of the 27-gene endocannabinoid signature. The first (23.9%) and second (14.3%) principal components were used. **(D)** COPD samples divided into female and male, and plotted separately as blue and orange-outlined box plots, respectively. For both (**A**) and (**C**) ellipses were added to represent 95% confidence intervals per sex. For both **(B)** and **(D)** log_2_-transformed expression values were plotted as box plots. The dashed line at zero represents the global baseline of expression for the entire set of genes.

Collectively, our data do not support sex differences in the endocannabinoid system in human airway epithelial cells from asthmatics or subjects with COPD.

### TRPV1 is up-regulated in airway epithelial cells from asthmatics

As our bioinformatic interrogation of the 27-gene endocannabinoid signature was restricted to genes, we next performed confirmatory protein expression analysis. The candidate we chose for validation was TRPV1, a confirmed receptor for cannabinoids that was the most differentially expressed candidate between our comparisons examining sex or disease status.

Using primary human airway epithelial cells from healthy donors or those with physician diagnosed asthma, protein was isolated from cells grown under submerged monolayer culture conditions. Immunoblot analysis confirms TRPV1 protein expression in human airway epithelial cells and revealed a qualitative increase in cells from asthmatics (**Figure 6**). TRPV1 staining normalized to total protein loading confirms a quantitative increase. In closing, our protein analysis is consistent with the bioinformatic analysis that revealed elevations in *TRPV1* gene in human airway epithelial cells from asthmatics.

**Figure 6:**
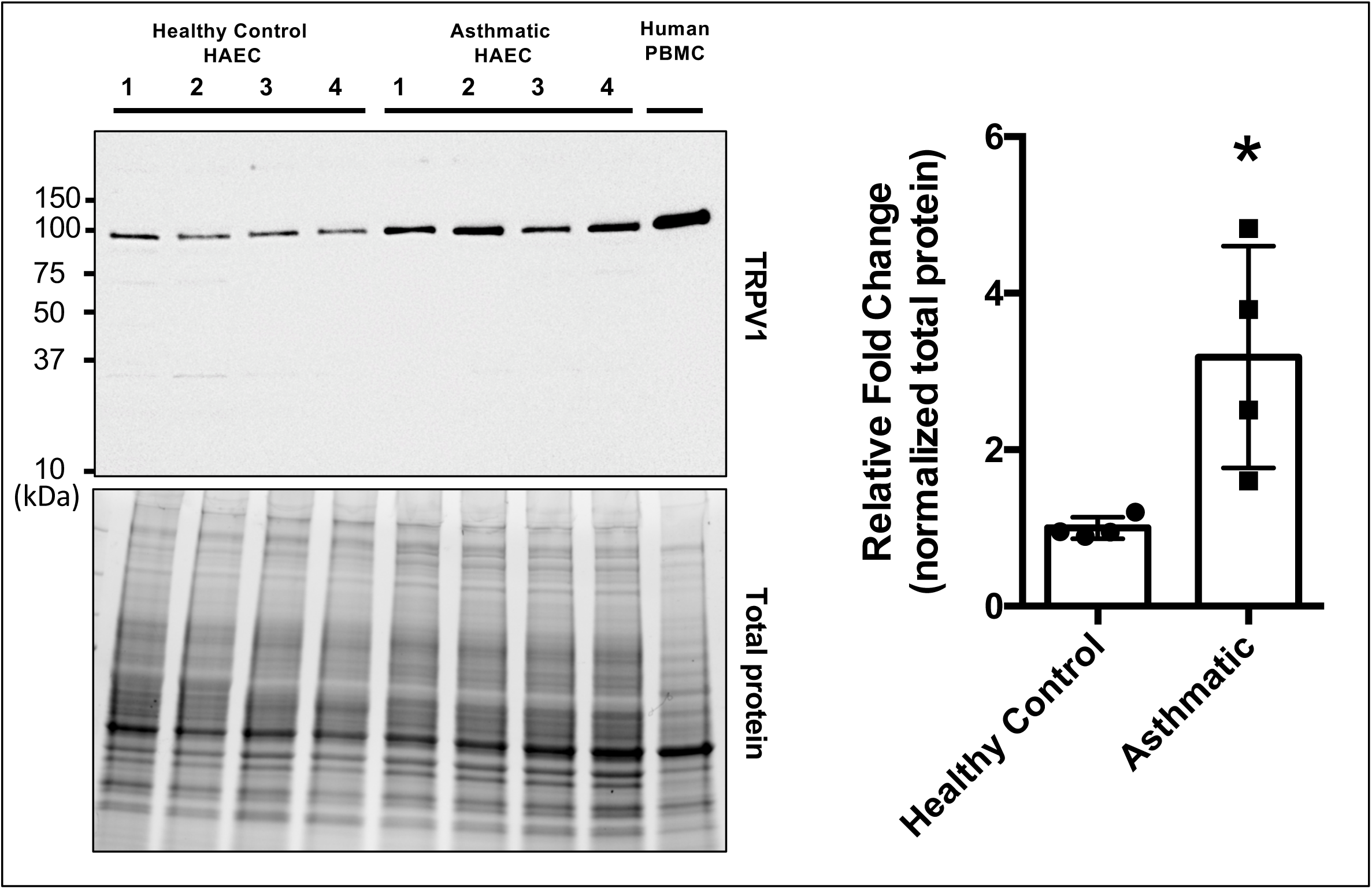
TRPV1 protein is elevated in human airway epithelial cells from asthmatics. Immunoblot of human airway epithelial cells (HAEC) from healthy subjects (n=4), asthmatics (n=4) and peripheral blood mononuclear cells (PBMC) (control) were analyzed for TRPV1 and quantified as fold change over healthy subjects, normalized to total protein loading. *=p<0.05.

## DISCUSSION

The dominant route of delivery of cannabis is via inhalation of combustion smoke, resulting in exposure to phytocannabinoids and activation of the endocannabinoid system(2-4). The airway epithelium represents the first line of defence in the human lung against inhaled insults including cannabis smoke. To better understand how the epithelium is able to respond to cannabis smoke exposure in the context of the endocannabinoid system, we performed a characterization study using bioinformatic and complementary protein analysis approaches. We demonstrate that three cannabinoid receptors, CB_1_, CB_2_, and TRPV1, are expressed at the protein level in human airway epithelium *in situ* and *in vitro*. The demonstration that these receptors were present warranted an exploration into the endocannabinoid system downstream of the receptors. Using 1090 unique patient samples of airway epithelial cells curated from publicly available datasets, we demonstrate in healthy subjects that the gene expression levels of the endocannabinoid system shows minor differences between females relative to males. In contrast, we demonstrate with samples from asthmatics or individuals with COPD that disease status appears to be a strong driver of endocannabinoid system gene expression, with no interaction with sex. Lastly, we validate the bioinformatic approach by demonstrating that TRPV1, a top candidate up-regulated at the gene level in our studies, is also up-regulated at the protein level in asthmatics. Collectively, our results confirm the expression of the endocannabinoid system in human airway epithelial cells and that disease status impacts expression to a greater extent then sex, which may have functional consequences in distinct populations of cannabis consumers.

The legalization of cannabis in multiple jurisdictions on a global scale reduces barriers for individuals to consume cannabis for either medicinal or recreational purposes. The dominant route of cannabis delivery is through inhalation of smoke from plant combustion(2-4). Inhaled cannabis smoke travels through the upper and lower airways, with airway epithelial cells being a major site of first contact. We have demonstrated with an *in vitro* model of airway epithelial cell culture that cannabis smoke induces a concentration-dependent reduction in airway epithelial cell viability, barrier function, while promoting pro-inflammatory cytokine secretion (31, 32). Complementary profiling of human epithelial cells isolated via bronchial brushings has demonstrated cannabis consumption-dependent elevations in TLR5, TLR6, and TLR9 gene expression(101). A limitation of these studies is the lack of mechanistic interrogation into the role that the endocannabinoid system contributed to the observed functional consequences of cannabis smoke. To begin to implicate the endocannabinoid system in epithelial cell functions, an *in vitro* approach has used direct cannabinoid administration to airway epithelial cells independent of combustion, showing functional consequences with altered barrier function mediated by mechanisms dependent and independent of cannabinoid receptors(35, 36). Our demonstration that multiple components of the endocannabinoid system are expressed at the gene level in airway epithelial cells with CB_1_, CB_2_, and TRPV1 receptors confirmed at the protein level are consistent with this system playing a contributing role in mediating cannabis smoke-induced effects. Collectively, our data and existing literature suggest that the dominant form of cannabis consumption, cannabis smoke inhalation, is able to induce functional responses in airway epithelial cells in a process that is mediated, at least in part, by the endocannabinoid system.

Cannabis use patterns have been reported to differ among males and females, with females consuming less in overall quantity and frequency(2-4). The endocannabinoid system is in turn regulated by sex hormones with diverse interactions at levels of receptors, enzymes, and signaling molecules (102, 103). In the context of lung health and disease, sex dependent lung physiology is observed as well as asthma and COPD disease incidence and progression (98, 99). In light of the potential for interactions between sex, user practices, and the endocannabinoid system, we performed a bioinformatic analysis that stratified for male and female sex status. In healthy individuals, we observe that only 6/27 of our endocannabinoid gene signature candidates were differentially regulated between males versus females (2 up-regulated, 4 down-regulated). In our analysis, the most significant differentially expressed gene between sexes was *ABHD6* (increased in female samples). Increased female expression of *ABHD6* has been previously reported in immune cells, with only female cells showing estrogen- or progesterone-dependent induction of *ABHD6* gene levels. Our observations of minor sex-dependent effects on the expression of the 27-gene endocannabinoid signature contrast with sex differences in the expression of CB_1_ and CB_2_ protein expression in heart tissue, where CB_1_ receptors are more highly expressed in females and CB_2_ receptors more highly expressed in males(94). Our observations of limited impact of sex on the 27-gene endocannabinoid signature expression in healthy subjects were conserved in asthmatics and individuals with COPD. In contrast to sex status, disease status drove a large effect for differential expression of the 27-gene endocannabinoid signature, with the majority of genes dysregulated (18/27 genes in asthmatics and 22/27 genes in subjects with COPD). Collectively, these findings suggest asthma or COPD status impacts expression of the endocannabinoid system, while sex status plays a comparatively smaller role.

The potential disease specific responses to cannabis exposure as a result of dysregulated endocannabinoid signaling is relevant as cannabinoids have and are being pursued for bronchodilatory and immunomodulatory properties (7, 8, 100, 104). Furthermore, in populations of asthmatics and COPD subjects, cannabis consumption is not avoided and shown to interact with disease progression. The reported benefits of cannabis exposure on lung function in asthmatics may be selective for this population based on endocannabinoid signaling. Indeed, if TRPV1 is a dominant receptor for responses downstream of inhaled cannabis, the signaling mediated in airway epithelial cells from asthmatics may be augmented relative to healthy controls. Our observation of elevated TRPV1 in airway epithelial cells from asthmatics is consistent with a previous report demonstrating correlation with asthma severity(105). A limitation of our study is that we did not explore other cell-types for TRPV1, which is also expressed in airway smooth muscle cells and can modulate smooth muscle contraction(106). In contrast, the lack of observed benefit on lung function and association with advanced pathology in COPD subjects may be a result of a distinct expression profile of the endocannabinoid system(10, 11, 100). Our results and those in the literature suggest that a universal response to inhaled cannabis by healthy subjects and those with asthma or COPD should not be assumed and cautions translation of safety and efficacy studies performed in healthy individuals to those with underlying asthma or COPD.

Our study is heavily focused on using deposited datasets generated by microarray gene expression technology. Microarrays are low cost, cover large numbers of transcripts, and benefit from a standardized format for analysis and public deposition of data. As a result of these benefits, gene expression data for large, independent cohorts are available for curation and data mining purposes if appropriate measures are taken to normalize and integrate datasets from diverse user groups. Despite benefits of curated microarray datasets, there exist limitations of our approach. Specifically, microarray technology is susceptible to poor signal to noise ratio, making transcripts expressed at low levels difficult to detect, suggesting that we may be under-estimating the signal or introducing high variance of these transcripts. To address this limitation, the SCAN method used functions to reduce technical and across-sample variation and to increase the signal to noise ratio while maintaining the ability to detect differentially expressed genes(87). As in all analyses, the sample size will dictate the statistical power, which can only be as large as the available datasets. In our case, the sample sizes for females and males in our asthma and COPD datasets were smaller than our healthy cohorts, which may have limited our ability to detect any sex-specific effects on endocannabinoid gene expression. Finally, in order to compare data across multiple experiments the microarray chip technology used needs to be considered with additional normalization methods required for data generated on different platforms. Different microarray technologies will have unique probe compositions and quality control protocols which can lead to systematic biases between experiments, though these batch-specific effects can be addressed using batch correction techniques and normalization(107). Despite these highlighted limitations, the curation of multiple microarray gene expression datasets from diverse cohorts of study subjects can be aligned with normalization methods to minimize batch and cross-platform effects and maximizing sample sizes to detect differential gene expression patterns.

In summary, we demonstrate that the endocannabinoid system is expressed in human airway epithelial cells at the gene level and CB_1_, CB_2_, and TRPV1 are expressed at the protein level. We demonstrate that gene expression patterns for a 27-gene endocannabinoid signature are differentially expressed between healthy individuals and those with asthma or COPD, whereas only minor differences are observed between sexes. We confirm that our bioinformatic approach for analysis of gene expression has the potential to reflect corresponding protein expression level changes as demonstrated by elevated TRPV1 protein expression in human airway epithelial cells from asthmatics relative to healthy controls. Our study lays a foundation with primary human lung samples from well defined patient populations to justify exploring the functional consequences of endocannabinoid system signaling in human airway epithelial cells in both health and disease. The complete functions of the endocannabinoid system in airway epithelial cells remain to be defined.

## ACKNOWLEDGEMENTS

The authors would like to acknowledge Olivia Marcello for her help in figure design.

