## Supplementary material for "Expression of the endocannabinoid system in the human airway epithelial cells – Impact of sex and chronic respiratory disease status": Table 1

| Molecule | Gene | Associated Protein | Function | Reference |
| --- | --- | --- | --- | --- |
| Receptor | *CNR1* | Cannabinoid receptor 1 (CB_1_) | Primary receptor involved in endocannabinoid signaling | Matsuda et al. 1990 |
|  | *CNR2* | Cannabinoid receptor 2 (CB_2_) | Primary receptor involved in endocannabinoid signaling | Munro et al. 1993 |
|  | *GABRA2* | Gamma-aminobutyric acid receptor subunit alpha-2 (GABRA2) | Implicated in cannabis dependency | Agrawal and Lynskey, 2010 |
|  | *GPR55* | G protein-coupled receptor 55 (GPR55) | Novel cannabinoid receptor | Ryberg et al. 2007 |
|  | *OPRM1* | Opioid receptor mu 1 (MOR1) | Implicated in cannabis dependency | Agrawal and Lynskey, 2010 |
|  | *TRPV1* | Transient receptor potential vanilloid 1 (TRPV1) | Novel cannabinoid receptor | Zygmunt et al. 1999 |
| Enzyme | *ABHD12* | 2-AG hydrolase ABHD12 (ABHD12) | Degradation of 2-AG | Blankman et al. 2009 |
|  | *ABHD6* | 2-AG hydrolase ABHD6 (ABHD6) | Degradation of 2-AG | Marrs et al. 2011 |
|  | *ADCY3* | Adenylyl cyclase 3 (AC) | Catalyzes formation of cAMP | Kano et al. 2009 |
|  | *AKT1* | AKT serine/threonine kinase 1 (AKT) | Regulates cell survival | Galve-Roperh et al. 2002 |
|  | *COMT* | Catechol-O-methyltransferase (COMT) | Degradation of dopamine | Henquet et al. 2006 |
|  | *CYP2C9* | Cytochrome P450 2C9 (CYP2C9) | Metabolism of THC | Watanabe et al. 2007 |
|  | *CYP3A4* | Cytochrome P450 3A4 (CYP3A4) | Metabolism of THC | Watanabe et al. 2007 |
|  | *DUSP6* | Dual specificity phosphatase 6 (MKP3) | Regulates MAPK signaling | Powles et al. 2004 |
|  | *FAAH* | Fatty acid amide hydrolase (FAAH) | Degradation of AEA | Kathuria et al. 2003 |
|  | *MAPK14* | Mitogen-activated protein kinase 14 (MAPK14) | Regulates cell survival | Onwuameze et al. 2013 |
|  | *MAP2K2* | Mitogen-activated protein kinase kinase 2 (MAP2K2) | Regulates cell survival | Powles et al. 2004 |
|  | *MAPK3* | Extracellular signal-regulated kinase (MAKP3) | Regulates cell survival | Bosier et al. 2008 |
|  | *MGLL* | Monoglyceride lipase (MAGL) | Degradation of 2-AG | Dinh et al. 2002 |
|  | *NOS2* | Inducible nitric oxide synthase (iNOS) | Inflammatory mediator | Cinar et al. 2016 |
|  | *PIK3CA* | Phosphatidylinositol-3-kinase (PI3K) | Regulates cell survival | Galve-Roperh et al. 2002 |
|  | *PRKACA* | Protein kinase A (PKA) | Regulates cell survival | Castillo et al. 2013 |
|  | *PTGS2* | Cyclooxygenase-2 (COX2) | Inflammatory mediator | Ruhaak et al. 2011 |
| Protein | *ABCB1* | P-glycoprotein 1 (P-GP) | Cannabinoid transportation | Kebir et al. 2018 |
|  | *GNAI1* | G_i/o_ alpha subunit (G_I/o_) | Coupled to cannabinoid receptors | Bondar and Lazar, 2017 |
|  | *NRG1* | Neuregulin 1 (NRG1) | Mediates cell-cell signaling | Hryhorowicz et al. 2018 |
|  | *TP53* | Tumor protein P53 (P53) | Regulates cell survival | Kim et al, 2012 |

Legend: AEA – anandamide, THC – tetrahydrocannabinol, 2-AG - 2-arachidonoylglycerol
