## Supplementary material for "Expression of the endocannabinoid system in the human airway epithelial cells – Impact of sex and chronic respiratory disease status": Table 2

| *GSE Accession* | *Affymetrix Chip* | *Asthma* | *COPD* | *Healthy* | *Citation* |
| --- | --- | --- | --- | --- | --- |
| GSE4302 | HG-U133 Plus 2 | 74 | 0 | 13 | Woodruff et al., 2007; Woodruff et al., 2009 |
| GSE4498 | HG-U133 Plus 2 | 0 | 0 | 12 (2F/10M) | Harvey et al., 2007; Tilley et al., 2009 |
| GSE5058 | HG-U133 Plus 2 | 0 | 14 (4F/10M) | 0 | Carolan et al., 2006; Tilley et al., 2009 |
| GSE7832 | HG-U133 Plus 2 | 0 | 0 | 8 (2F/6M) | Tilley et al., 2009 |
| GSE8545 | HG-U133 Plus 2 | 0 | 15 (3F/12M) | 5 (1F/4M) | Ammous et al., 2008 |
| GSE10006 | HG-U133 Plus 2 | 0 | 20 (4F/16M) | 21 (2F/19M) | Carolan et al., 2008 |
| GSE11784 | HG-U133 Plus 2 | 0 | 17 (1F/3M) | 40 (18F/22M) | Tilley et al., 2011 |
| GSE11906 | HG-U133 Plus 2 | 0 | 0 | 30 (10F/20M) | Raman et al., 2009 |
| GSE13931 | HG-U133 Plus 2 | 0 | 0 | 19 (4F/15M) | Carolan et al., 2009 |
| GSE13933 | HG-U133 Plus 2 | 0 | 0 | 11 (6F/5M) | Turetz et al., 2009 |
| GSE14224 | HuEx-1.0-st-v2 | 0 | 0 | 11 (7F/4M) | Schembri et al., 2009 |
| GSE17905 | HG-U133 Plus 2 | 0 | 0 | 1 (1M) | Wang et al., 2010 |
| GSE19667 | HG-U133 Plus 2 | 0 | 0 | 3 (3F) | Strulovici-Barel et al., 2010 |
| GSE20257 | HG-U133 Plus 2 | 0 | 1 (1F) | 0 | Shaykhiev et al., 2011 |
| GSE22047 | HG-U133 Plus 2 | 0 | 23 | 81 | Butler et al., 2011 |
| GSE34450 | HG-U133 Plus 2 | 0 | 0 | 11 | Wang et al., 2012 |
| GSE37147 | HuGene-1.0-st-v1 | 0 | 110 (35F/52M) | 8 | Steiling et al., 2013 |
| GSE40364 | HG-U133 Plus 2 | 0 | 7 | 0 | Gao et al., 2014 |
| GSE43079 | HG-U133 Plus 2 | 0 | 0 | 16 | Buro-Auriemma et al., 2013 |
| GSE43939 | HG-U133 Plus 2 | 0 | 0 | 13 | Hessel et al., 2014 |
| GSE52237 | HG-U133 Plus 2 | 0 | 0 | 2 | Walters et al., 2014 |
| GSE64614 | HG-U133 Plus 2 | 0 | 0 | 31 | Yang et al., 2017 |
| GSE67472 | HG-U133 Plus 2 | 62 (34F/28M) | 0 | 43 (20F/23M) | Christenson et al., 2015 |
| GSE77658 | HG-U133 Plus 2 | 0 | 0 | 6 | Zhou et al., 2016 |
| GSE84101 | HG-U133 Plus 2 | 0 | 0 | 7 | Walters et al., 2017 |
| GSE97010 | HuGene-1.0-st-v1 | 0 | 0 | 126 (28F/98M) | Billatos et al., 2018 |
| GSE108134 | HG-U133 Plus 2 | 0 | 131 | 98 | O’Beirne et al., 2018 |
